# La1: an evolutionarily conserved player in the Arabidopsis telomerase complex

**DOI:** 10.64898/2026.01.25.701547

**Authors:** Chinmay Phadke, Saundarya Mishra, Jiarui Song, Rebekah Holtsclaw, Claudia Castillo Gonzalez, Ishan Kundel, Edward M. Marcotte, Ophelia Papoulas, Dorothy E. Shippen

## Abstract

Analysis of yeast and mammalian telomerase ribonucleoprotein (RNP) complexes reveals striking divergence in their biogenesis and protein complements. However, little is known about telomerase in plants. In addition to the catalytic subunit TERT and the templating RNA TR, we previously reported *Arabidopsis thaliana* contains two telomerase accessory factors, AtNAP57, a dyskerin homolog that in mammals is essential for telomerase activity, and the telomeric DNA binding protein AtPOT1a. Both proteins stimulate Arabidopsis telomerase repeat addition processivity. Here we employ quantitative mass spectrometry (MS) to further examine telomerase in Arabidopsis. Unexpectedly, dyskerin and AtPOT1a were not detected in our purified complexes, but AtLa1, an RNA-binding factor that recognizes the UUU-3′OH of RNA Pol III transcripts, was highly enriched. RNA-IP assays confirmed AtLa1 association with AtTR in vivo. RNAi-mediated knockdown of AtLa1 strongly diminished telomerase activity, indicating AtLa1 is required for its function in vivo. In vitro binding studies revealed that AtLa1 contacts AtTR via the UUU-3′OH and a plant-specific P1a-P1b-P4 three-way junction (TWJ). Since the TWJ is also required for AtNAP57 binding, the data suggesting that AtNAP57 and AtLa1 compete for AtTR binding or sequentially associate during RNP during biogenesis. In contrast to AtNAP57, AtLa1 did not stimulate telomerase activity when TERT and TR were assembled in vitro, consistent with function during a different step in telomerase assembly. We conclude Arabidopsis telomerase employs multiple accessory factors utilized by both mammalian and single-celled relatives. Further exploration of Arabidopsis telomerase may offer novel insight into telomerase evolution and mechanisms of biogenesis.

**Significance of Research:** Quantitative mass spectrometry of Arabidopsis telomerase uncovered AtLa1, a homolog of ciliate and yeast proteins that promotes telomerase maturation. AtLa1 is essential for telomerase function in vivo, and in vitro it engages the same region of AtTR bound by AtNAP57, homologous to a telomerase accessory from mammals.

## Introduction

Telomeres consist of repetitive G-rich DNA sequences that cap chromosome ends. Telomeric DNA is synthesized by the telomerase reverse transcriptase, a ribonucleoprotein (RNP) enzyme that contains two core subunits: a catalytic protein, Telomerase Reverse Transcriptase (TERT), and an RNA moiety with a telomere templating sequence termed telomerase RNA (TR). Although TERT and TR are sufficient to reconstitute telomerase activity in vitro, an array of accessory factors assemble with the core in vivo to facilitate RNP maturation and enzyme engagement and regulation at chromosome termini (Podlevsky and Chen, 2016; Davis et al., 2023; Davis and Chakrabarti, 2024). The protein composition of telomerase varies widely across eukaryotic lineages. Studies of mammalian, yeast and ciliate complexes reveal a multitude of protein-protein and RNA-protein interactions are required to form functional telomerase holoenzyme (Davis and Chakrabarti, 2024). Some proteins stably associate with the core, while others interact transiently during RNP biogenesis and maturation.

Human telomerase harbors a 451-nt telomerase RNA (hTR) transcribed by RNA polymerase II (Feng et al., 1995; Cong et al., 1999). Biochemical and cryo-EM studies reveal that the holoenzyme carries two copies of NHP2, NOP10, TCAB1, GAR1 and dyskerin, which together form a conserved H/ACA RNP scaffold essential for hTR stability and telomerase function (Chen et al., 2000; Venteicher et al., 2009; Nguyen et al., 2018; Ghanim et al., 2021). These proteins assist in RNA folding and facilitate its maturation and assembly into an active enzyme. They also promote proper localization of the telomerase complex within the nucleus, ensuring it reaches sites where telomere elongation occurs. Of particular note, dyskerin interacts with an H/ACA box near the 3’ terminus of hTR, contributing to RNA stability and maturation (Nguyen et al., 2018; Ghanim et al., 2021).

The budding yeast telomerase features a different set of protein components, including Est1, Est3, Pop1/6/7, yKu and the Sm□ ring complex. These accessory proteins support the stability, maturation and activity of the *Saccharomyces cerevisiae* telomerase RNA (TLC1) and the telomerase enzyme complex in vivo. Est1 and Est3 are telomerase-specific regulatory proteins that recruit telomerase to telomeres and modulate its activity, with Est1 bridging the interaction between TLC1 and the telomere-binding protein Cdc13, and Est3 associating in a cell cycle–dependent manner (Lendvay et al., 1996; Chan et al., 2008; Chen et al., 2018). The Pop1/6/7 complex, shared with RNase P/MRP, binds to a conserved region of TLC1 and stabilizes the RNA, enhancing association with Est1 and Est2 (Lemieux et al., 2016; Garcia et al., 2020). The yKu heterodimer (yKu70/yKu80) binds to a stem-loop structure in TLC1 and is essential for its nuclear retention and telomeric localization, particularly during G1 (Fisher et al., 2004; Hass and Zappulla, 2015; Garcia et al., 2020). The Sm□ ring complex, a conserved group of RNA-binding proteins, associates with the 3′ end of TLC1 and plays a key role in its processing, stability and nuclear import (Seto et al., 1999; Vasianovich et al., 2020).

In *Schizosaccharomyces pombe*, a different set of proteins engage the telomerase RNA, including the La-related protein (LARP) Pof8, which promotes enzyme activity (Páez-Moscoso et al., 2018; Davis and Chakrabarti, 2024). This finding is curious since La proteins typically associate with transcripts synthesized by RNA Pol III, but both the budding and fission yeast telomerase RNA molecules are generated by RNP Pol II. In addition, the majority of telomerase accessory factors are specific to yeast and not found in higher eukaryotes, emphasizing lineage-specific adaptations in telomerase biogenesis and regulation (Zapulla and Cech, 2004; Lemieux et al., 2016).

The ciliate *Tetrahymena thermophila* possesses yet another structurally and compositionally distinct telomerase holoenzyme. TR is transcribed by RNA Pol III, and accessory proteins such as p65, p75, p45, p19, p50, Teb1, Teb2 and Teb3 associate with the core enzyme (Jiang et al., 2013; Nguyen et al., 2018). Teb1 is a single-stranded DNA-binding protein that enhances telomerase processivity, functionally analogous (though not homologous) to human POT1 (Min and Collins, 2010). Other subunits, such as p75 and p45, bridge the core RNP with the DNA substrate, supporting both recruitment and catalytic activity. Relevant to this study, p65, a LARP unique to Tetrahymena (Akiyama et al., 2012; Singh et al., 2012) promotes proper TR folding and stability.

Telomerases from the plant kingdom are less understood. Only recently was the bona-fide TR identified (Fajkus et al., 2019; Song et al., 2021). Like other telomerase RNAs, the *Arabidopsis thaliana* RNA (AtTR) contains conserved core structural and functional domains, including a template region that guides telomere repeat synthesis and a pseudoknot domain that contributes to the structural integrity of TR and its catalytic function (Fajkus et al., 2019; Song et al., 2019; Štefanovie et al., 2024). AtTR also displays plant-specific features, including a unique three-way junction (TWJ) and a distinctive organization of hairpin loops and stem regions not found in vertebrate or yeast counterparts (Song et al., 2019; Štefanovie et al., 2024). These structural folds likely reflect plant-specific modes of RNA processing, stabilization and holoenzyme assembly.

A multitude of putative accessory proteins have been reported for Arabidopsis telomerase (Schrumpfová et al., 2014), but the functional relevance of most of these factors in telomere biology is unclear. For example, Telomere Repeat Binding (TRB) proteins from *A. thaliana* interact with both AtTERT and AtPOT1b, suggesting roles in telomere protection and chromatin organization (Dvořáčková et al., 2015). TRB4 and TRB5 also function as transcriptional activators of Polycomb Repressive Complex 2 (PRC2)-regulated genes, implying a connection between telomere biology and developmental processes such as flowering time control (Kusová et al., 2023). Several additional telomerase-associated proteins, including some involved in chromatin remodeling and transcriptional regulation, have been identified through yeast two-hybrid screening and co-immunoprecipitation assays. These factors include chromatin-associated proteins such as MSI1, a histone-binding protein linked to PRC2 function; NRP1 and NRP2, which are involved in nucleosome assembly; and VIM1, associated with DNA methylation (Schrumpfová et al., 2014; Fulnečková et al., 2021). Although these findings underscore the multifunctionality of telomerase-associated proteins in coordinating chromosome end maintenance with broader epigenetic and developmental programs in plants, they provide limited mechanistic insight into telomerase RNP assembly, enzymatic function and evolution.

We previously reported that *A. thaliana* telomerase engages AtPOT1a in vivo and in vitro (Surovtseva et al., 2007; Arora et al., 2016). Protection of Telomeres (POT1) proteins in yeast and in mammals are stable components of the telomere and protect against a DNA damage response (Baumann and Cech, 2001; Hockemeyer et al., 2005). By contrast, AtPOT1a accumulates at telomeres during S-phase, and loss of AtPOT1a results in progressive telomere shortening, similar to loss of TERT (Surovtseva et al., 2007). AtPOT1a is associated with enzymatically active telomerase, binding TERT independently of TR and stimulating repeat addition processivity (Surovtseva et al., 2007; Renfrew et al. 2014; Song et al., 2021). The dyskerin homolog (AtNAP57) also interacts with Arabidopsis telomerase (Kannan et al., 2008; Song et al., 2021) as well as telomerase from *Allium cepa* (Fajkus et al., 2019). Analysis of an AtNAP57 loss-of-function mutant showed reduced telomerase activity and a shorter telomere length set point, indicating that dyskerin is essential for telomere homeostasis in Arabidopsis (Kannan et al., 2008). Unlike AtPOT1a, AtNAP57 interacts with telomerase in an RNA-dependent manner (Song et al., 2021), similar to dyskerin in vertebrate telomerase (Mitchell et al., 1999; Garus and Autexier, 2021). Notably, all the plant TR molecules characterized to date are transcribed by RNA Pol III (Fajkus et al., 2021). Therefore, the interaction of AtNAP57 with AtTR is unexpected. Biochemical studies revealed that AtNAP57 binds the TWJ within AtTR (Song et al., 2021), arguing that a plant-specific adaptation facilitates this protein-RNA interaction.

In this study, we employ quantitative mass spectrometry to identify additional proteins that associate with *A. thaliana* telomerase. We present AtLa1 as a novel and essential telomerase-interacting protein and show that AtLa1 engages AtTR through the same domains as dyskerin. Our findings argue that AtLa1 and dyskerin are involved in a sequential assembly process during the biogenesis and maturation of Arabidopsis telomerase.

## Results

### Quantitative mass spectrometry reveals a novel telomerase-interacting protein, AtLa1

To identify proteins associated with the *A. thaliana* telomerase core complex, we prepared a transgenic ‘super-telomerase’ line that overexpresses both AtTERT and AtTR by employing a binary vector carrying a 35S promoter-driven TwinStrep (TSII) tagged AtTERT and a U6 promoter-driven AtTR. Tagged-AtTERT was cloned using the genomic AtTERT sequence (5380 bp), including all introns with a TSII tag attached to the N-terminus, and the AtTR sequence (498 bp) was cloned along with the U6 promoter and terminator. This binary vector was transformed into *tert* null mutants. We identified two independent lines, 12-5 and 10-3, which exhibited 2-fold and 40-fold higher telomerase activity, respectively, than the wild type (Barcenilla et al., 2023). Terminal Restriction Fragment (TRF) analysis indicated these lines complemented the *tert* null allele and exhibited slightly longer telomeres than wild type (Barcenilla et al., 2023). We extracted protein from inflorescences of line 10-3, as well as *tert* mutant controls, and the TSII-tagged AtTERT was captured on strep-tactin beads. The eluted proteins were concentrated and subjected to quantitative mass spectrometry analysis (Fig. 1A).

**Fig. 1.**
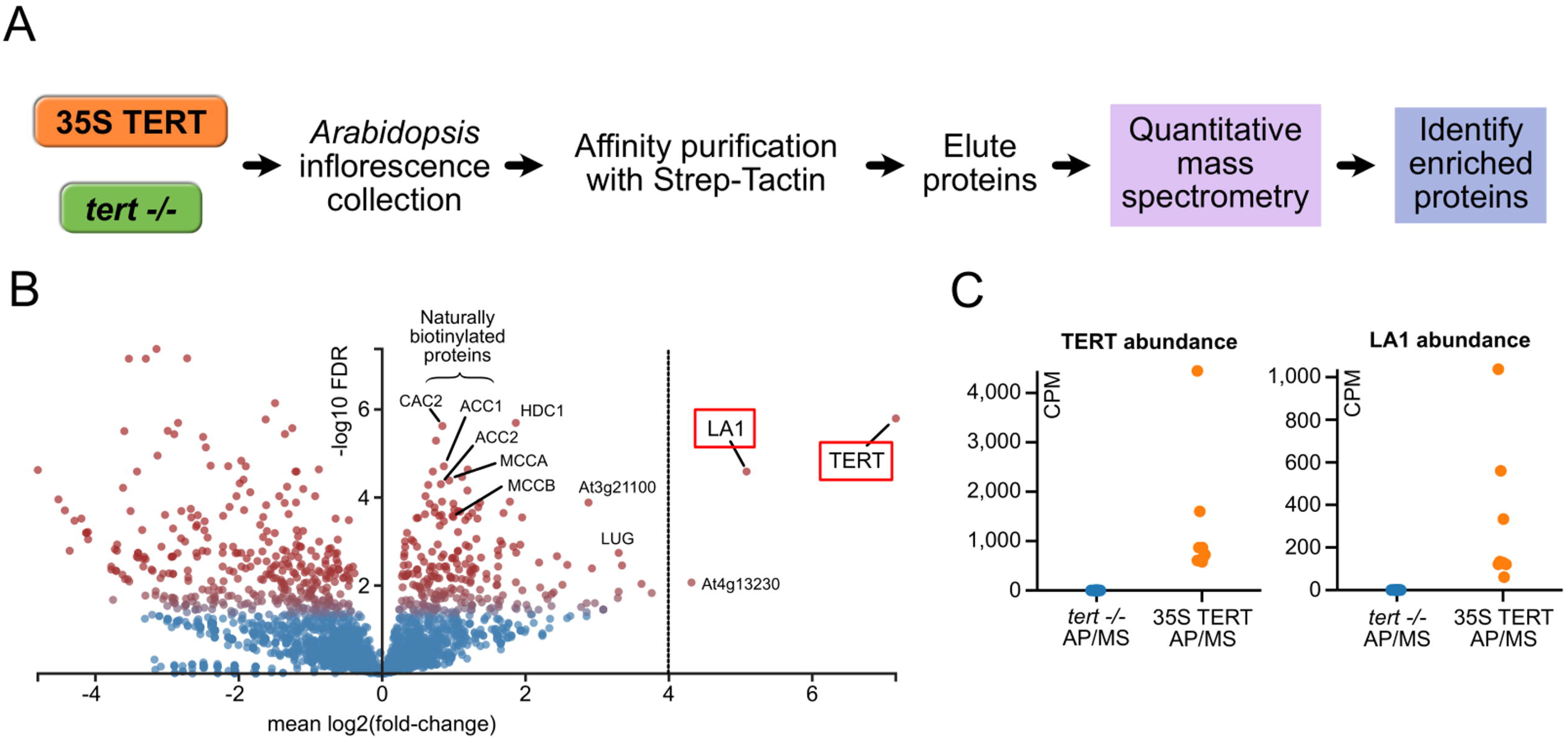
AtLa1 co-purifies with *A. thaliana* telomerase RNP. (A) Protocol for the extraction and affinity purification of telomerase RNP. (B) Volcano plot of protein fold-change (x-axis) and statistical significance (y-axis; FDR, false discovery rate, as measured by EdgeR) in affinity purifications from 35S TERT plants relative to the untransformed control plants (*tert −/−*), showing that AtLa1 is the most highly enriched protein after TERT and hence likely involved in the telomerase RNP. AtTERT and AtLa1 are highlighted in red. (C) Mass spectrometry-measured protein abundances (in counts per million, cpm) for AtTERT and AtLa1, showing the high level of enrichment compared to *tert* null controls.

Relative protein abundances in super-telomerase plants compared to the *tert* mutant plants were quantified from four biological replicates, using label-free protein quantification. Differentially enriched proteins were identified (see Methods) and are displayed in Fig. 1B as a volcano plot where both axes are log-transformed, enabling visualization of proteins statistically enriched in the affinity-purified samples. As expected, AtTERT was the most highly enriched protein in super-telomerase relative to the *tert* mutant samples, with an enrichment significantly greater than the naturally biotinylated proteins observed in this analysis, which might be expected to be at comparable levels in both the super-telomerase and *tert* mutant samples. Interestingly, the next most significantly enriched protein was the La-related RNA-binding protein AtLa1 (Fleurdépine et al., 2007), with a greater than 30-fold enrichment relative to the *tert* null control, which showed negligible background (Fig. 1C). Since La-related proteins have been detected in multiple different telomerase complexes (Akiyama et al., 2012; Singh et al., 2012; Paez-Moscoso et al., 2018; Davis and Chakrabarti, 2024), we assessed the functional relevance of this interaction.

### AtLa1 is required for telomerase function *in vivo*

AtLa1 is a bona-fide RNA-binding protein of Arabidopsis, essential for embryonic development (Cui et al., 2015) and it maintains stem cell homeostasis by initiating translation of WUSCHEL (WUS) mRNA (Cui et al., 2015). It is primarily localized in the nucleoplasm; environmental hazard promotes its nucleocytoplasmic translocation enhancing WUSCHEL (WUS) mRNA translation (Cui et al., 2015). AtLa1 binds the 3’ UUU of Pol III transcripts (Fleurdépine et al., 2007), and the canonical RNA recognition motifs (RRM) of AtLa1 are sufficient to restore La protein function in yeast, including U6 snRNA biogenesis, implying a conserved role in RNP assembly.

To investigate the biological relevance of AtLa1 for Arabidopsis telomerase, we used a previously characterized transgenic line of *A. thaliana* overexpressing tagged AtLa1 (35S:HA:AtLa1_syn_) (Cui et al., 2015). Quantitative PCR and western blotting verified over-expression of tagged AtLa1 relative to wild type Col-0 (Fig. 2A, 2B and 2C). Specifically, we detected 50-to-150-fold higher transcript levels and 8.9-to-9.5-fold higher protein expression in 35S:HA:AtLa1_syn_ compared to the untransformed wild type Col-0 control.

**Fig. 2.**
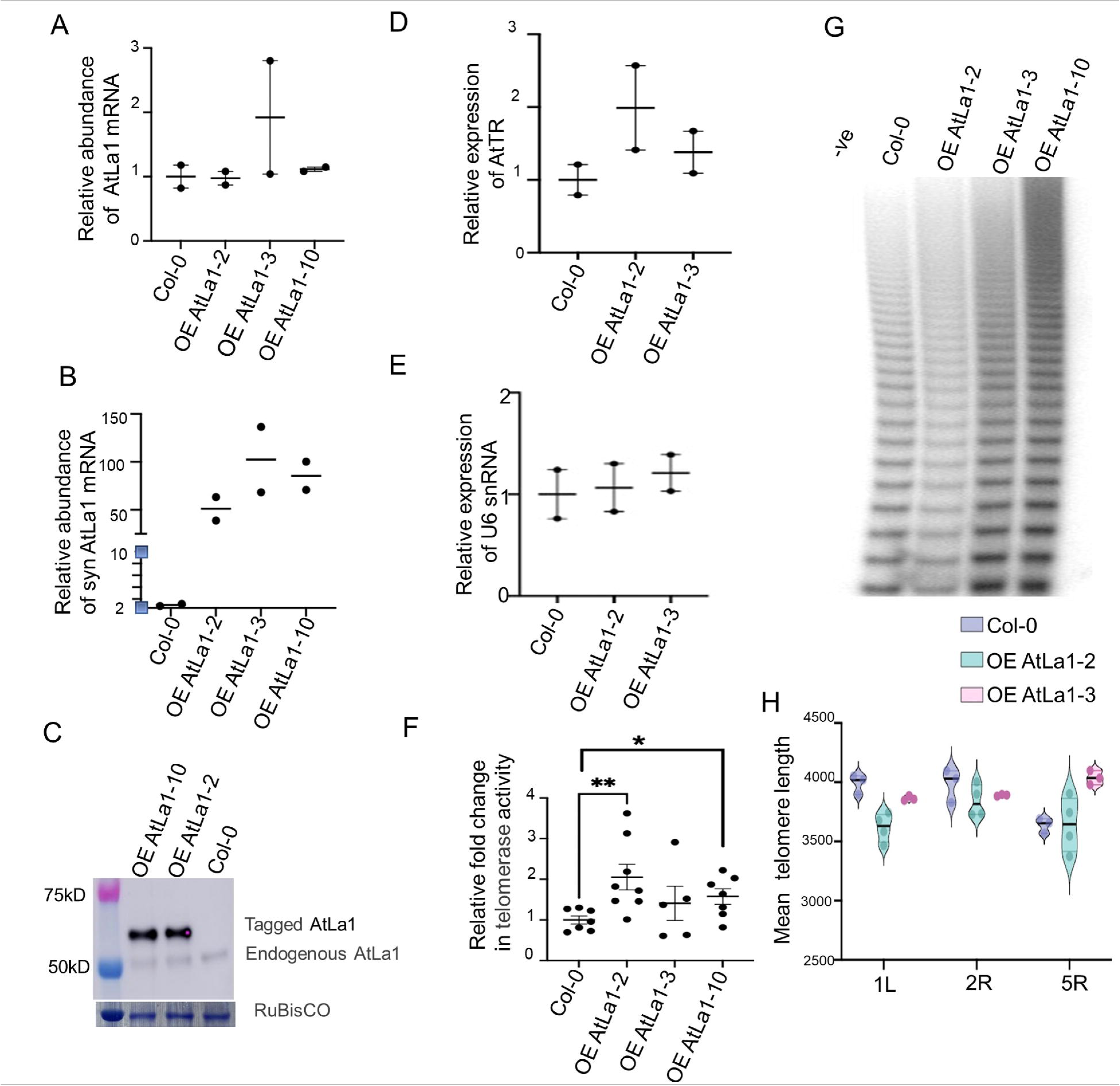
Over-expression of AtLa1 increases telomerase activity in vivo. Characterization of AtLa1 over-expression lines with wild type Col-0, (A) Quantitative PCR analysis for endogenous AtLa1 transcripts suggests mild change in transcripts and data normalized with *ACTIN2* and (B) Quantitative PCR results for AtLa1 syn transcripts normalized with *ACTIN2*. **(C)** Western blotting results for tagged and endogenous AtLa1 protein expression in transformed lines OE AtLa1-10, OE AtLa1-2 as compared to wild type Col-0. Blots probed with α-AtLa1 primary antibody, and it suggests 10-fold higher protein expression as compared to Col-0 and RuBisCO acts as a loading control. (D and E) Quantitative PCR analysis of AtTR and U6 snRNA transcript abundance. It is normalized with GAPDH. (F and G) Results for Quantitative Telomeric Repeat Amplification Protocol (qTRAP) and conventional gel-based TRAP assays of lines over-expressing AtLa1. (H) Mean telomere length of PETRA data analyzed by WALTER.

We also monitored the steady state level of AtTR transcripts in plants over-expressing AtLa1, reasoning that if AtLa1 is important for AtTR stability, AtTR levels might increase. We found a modest (1.5-fold) increase in AtTR and but no increase in U6 snRNA transcripts in the 35S:HA:AtLa1_syn_ lines compared to wild type (Fig. 2D and 2E). We measured telomerase enzyme activity using conventional TRAP and quantitative TRAP (qTRAP) to check how AtLa1 over-expression affects telomerase activity. We found a 2.5 to 3.5-fold increase in telomerase activity in the transgenic lines (Fig. 2F and G), implying that AtLa1 promotes telomerase activity. Telomere length analysis using primer extension telomere repeat amplification (PETRA) revealed no significant change when AtLa1 was over-expressed (Fig. 2H, S1A, S1B, S1C), consistent with previous studies showing that elevated levels of telomerase enzyme activity do not necessarily lead to telomere elongation in Arabidopsis (Barcenilla et al., 2023).

Since AtLa1 is required for embryogenesis (Fleurdépine et al., 2007), we examined the impact of AtLa1 depletion using an inducible RNAi knock-down line. Specifically, we created a β-estradiol-inducible RNAi knock-down construct to control AtLa1 expression post-embryonically. We also created stable homozygous T3 transformed lines containing the pER8-hpRNAiAtLa1 knock-down construct. These lines were considered in parallel with wild type Col-0 and were plated on 0.5 MS supplemented with 25μM/ml β-estradiol (Fig. 3A). Western blotting with 7-day-old seedlings showed a 10 to 60-fold reduction in AtLa1 protein expression in 4-3-1 and 1-7-6 lines, respectively, as compared to the control (Fig. 3B). We also asked how AtLa1 depletion impacted AtTR and U6 snRNA accumulation in the knock-down lines. We found no significant decrease in either RNA relative to the GAPDH control (Fig. 3C and D).

**Fig. 3.**
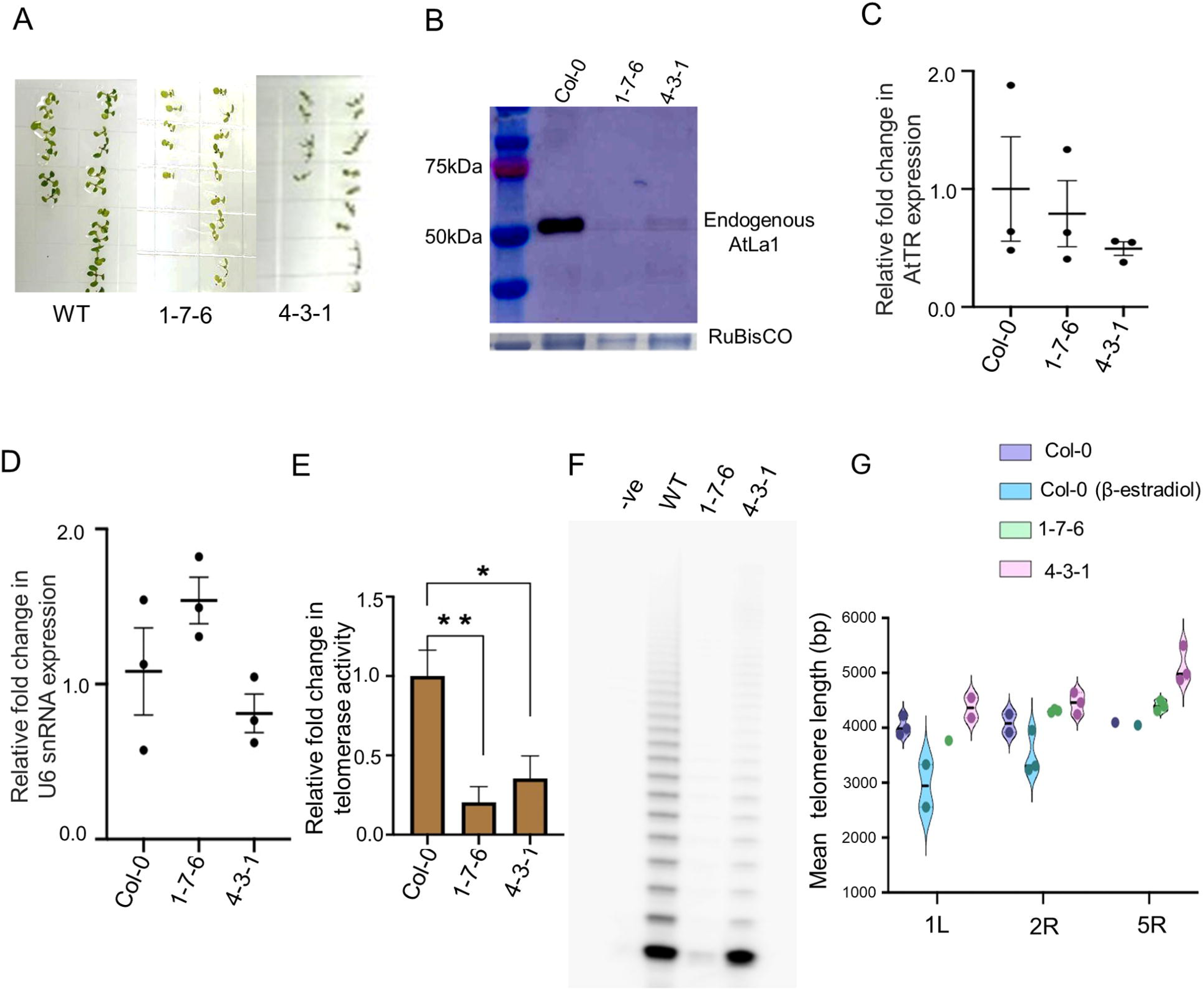
Knock-down of AtLa1 reduces telomerase activity. (A) Reduced growth observed in seven-day-old seedlings transformed with an inducible RNAi construct (lines 1-7-6 and 4-3-1) relative to the wild type (Col-0). Seedlings were grown in MS plates supplemented with 25 µM β-estradiol. (B) Characterization of inducible RNAi knockdown lines by Western blot analysis (blot probed with anti-AtLa1 primary antibody). RuBisCO served as a loading control. (C and D) Results qPCR analysis for AtTR and U6 snRNA abundance in knockdown transformed lines as compared to Col-0. Col-0 and knockdown lines data were normalized with GAPDH. (E and F) Results of Telomerase activity assays (qTRAP and TRAP) for AtLa1 inducible RNAi knockdown lines (1-7-6 and 4-3-1) as compared to wild type Col-0. (G) Average telomere length analyzed in Col-0, Col-0(β-estradiol) and incubated RNAi knockdown lines 1-7-6 and 4-3-1by WALTER analysis.

We unexpectedly observed a decrease in telomerase activity in the 25μM/ml β-estradiol treated wild type Col-0 control as compared to untreated Col-0 (Fig. S2A). Therefore, we focused our analysis of telomerase on stable transformed lines. Telomerase activity measured by qTRAP decreased by six-fold in the induced 1-7-6 line and four-fold in the 4-3-1 line, relative to wild type Col-0 control (Fig. 3E). We also examined the profile of telomere elongation products created in these lines using a standard gel-based conventional TRAP assay. In one experiment we observed faint backgrounds of elongation products in negative control, but much less than in the wild type Col-0 sample (Fig. S2B). Importantly, the abundance of products obtained for the knock-down lines was similar to the negative control (Fig 3F, S2B). In addition, while the overall intensity of the product bands was fainter for the knock-down lines, the number telomere repeats added (the length of the products) was approximately the same, indicating that loss of AtLa1 did not specifically impact the repeat addition processivity of telomerase.

Analysis of telomere length in plants seven days post AtLa1 knock-down revealed no significant change in telomere length relative to the non-transformed control lines (Fig. 3G, S2C, S2D and S2E). Nevertheless, the striking reduction in telomerase enzyme activity when AtLa1 levels are decreased argues that AtLa1 is required for telomerase function in *A. thaliana*.

### AtLa1 binds AtTR via a plant-specific TWJ and the UUU-3’OH

To test whether AtLa1 is associated with AtTR *in vivo*, we used 35S:HA:AtLa1_syn_ for RNA-IP. Western blotting confirmed that HA-tagged AtLa1 was enriched when extracts from plants expressing 35S:HA:AtLa1_syn_ were subjected to affinity purification (Fig. 4A). We used qPCR to determine if AtTR was likewise enriched in this purification. U6 snRNA was employed as a positive control since AtLa1 has been shown to associate with this RNA (Fleurdépine et al., 2007). Relative to the untransformed wild type control which showed no enrichment, the abundance of AtTR and U6 snRNA was increased by 2 and 2.5-fold, respectively (Fig. 4B). These findings provide additional support that AtLa1 is specifically associated with Arabidopsis telomerase.

**Fig. 4.**
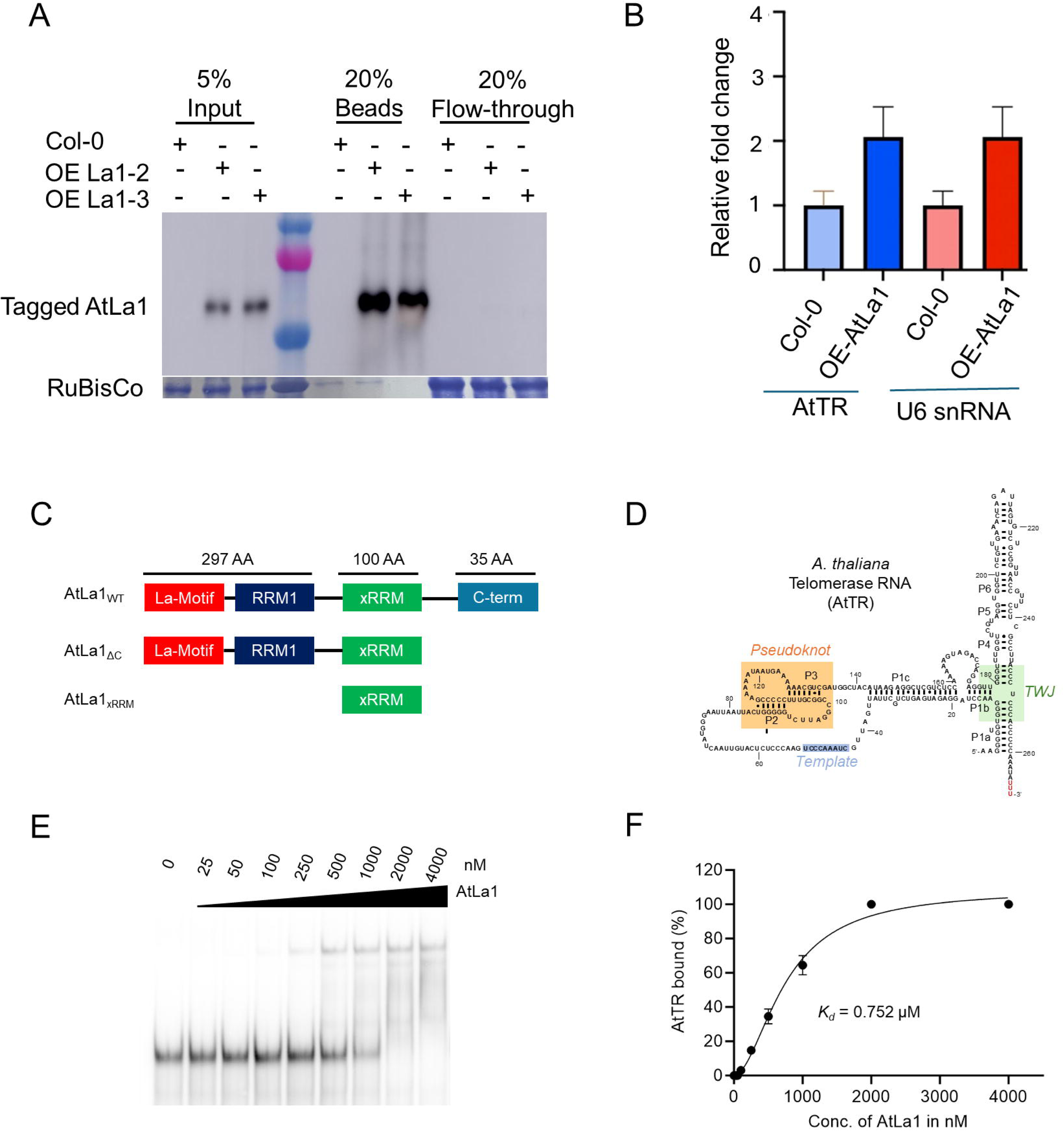
AtLa1_WT_ binds full-length AtTR in vitro. (A and B) RNA immunoprecipitation (RIP) suggests interaction of AtLa1 interact with AtTR in vivo condition confirmed by western blot (probed with anti-HA-HRP) and Results of RIP. (C) Domain organization of AtLa1_WT_ (433 amino acids) and its truncations used in EMSA. Radiolabeled AtTR was incubated with increasing concentrations of recombinant AtLa1 protein (0–4 μM). (D) Secondary structure schematic of *Arabidopsis thaliana* telomerase RNA (AtTR), highlighting the pseudoknot (PK), template domain and plant-specific three-way junction (TWJ). (E) Electrophoretic mobility shift assay (EMSA) showing binding of purified AtLa1_WT_ to in vitro transcribed and PAGE-purified AtTR. (F) Quantification of EMSA data reveals that AtLa1_WT_ binds AtTR with a dissociation constant (*K_d_*) of 0.752 µM.

Since our RNA-IP data indicated that AtLa1 is associated with AtTR, we used Electrophoretic Mobility Shift Assays (EMSA) to investigate whether AtLa1 binds AtTR directly. We first tested AtTR interaction using full-length AtLa1 (AtLa1_WT_) (433 amino acids: 6xHis-SUMO-AtLa1_WT_) (Fig. 4C). AtLa1_WT_ was expressed and purified from *E. coli* (Rosetta cells) (Fig. S3). Fractions 13 and 14 with a purity of ∼95% were used for the EMSA assay. Full-length wild type AtTR was prepared by *in vitro* transcription, and the RNA was purified by 6% denaturing PAGE (Fig. 4D). EMSA data indicated that AtLa1_WT_ binds AtTR *in vitro* with a dissociation constant (*K_d_*) of 0.752 µM (Fig. 4E and 4F).

AtLa1 contains a canonical La domain, followed by RNA Recognition Motif 1 (RRM1), a poorly characterized RRM2 (also referred to as xRRM) and a short C-terminal region (Fig. 4C) (Bousquet-Antonelli and Deragon, 2009). To dissect the contribution of individual domains in AtLa1 toward AtTR binding, we expressed and purified two truncation constructs in *E. coli*: 6xHis-SUMO–AtLa1_ΔC_, lacking the 36-residue C-terminal region; and 6xHis-SUMO–AtLa1_RRM2_, a 100-residue construct comprising only RRM2 (Fig. S3A and S3C). EMSAs revealed that neither of the two truncation constructs was capable of binding AtTR (Fig. S4A and S4B). These findings suggest that the canonical La domain, RRM1, as well as the C-terminal region of AtLa1 are needed for AtTR interaction.

To pinpoint the binding site of AtLa1 on AtTR, we prepared several truncation constructs of AtTR (Fig. 5 and 6) and studied their binding with AtLa1_WT_. We first asked if the core structural and functional domains within AtTR are necessary for AtLa1 binding. Truncations A and B lack the pseudoknot (PK) and template region, while truncation B also lacks the bulge between P1c and P1b. We found that AtLa1_WT_ retains binding to truncations A and B, but relative to full-length AtTR with approximately three-fold lower affinity (*K_d_* =2.40 µM) and approximately two-fold lower affinity with truncation B (*K_d_* = 1.62 µM), respectively (Fig. 5A and 5B). Thus, the PK and template domains of AtTR are not essential for AtLa1 binding. To further investigate how these regions contribute to AtLa1 engagement, we prepared truncations G and H, which retain PK and template regions but lack the 3’ stem P1a. Truncation H also lacks P4, P5, and P6. Interestingly, no binding was detected for truncations G or H, suggesting that some feature of the 3’ stem region is needed for interaction with AtLa1_WT_ (Fig. 5C and 5D).

**Fig. 5.**
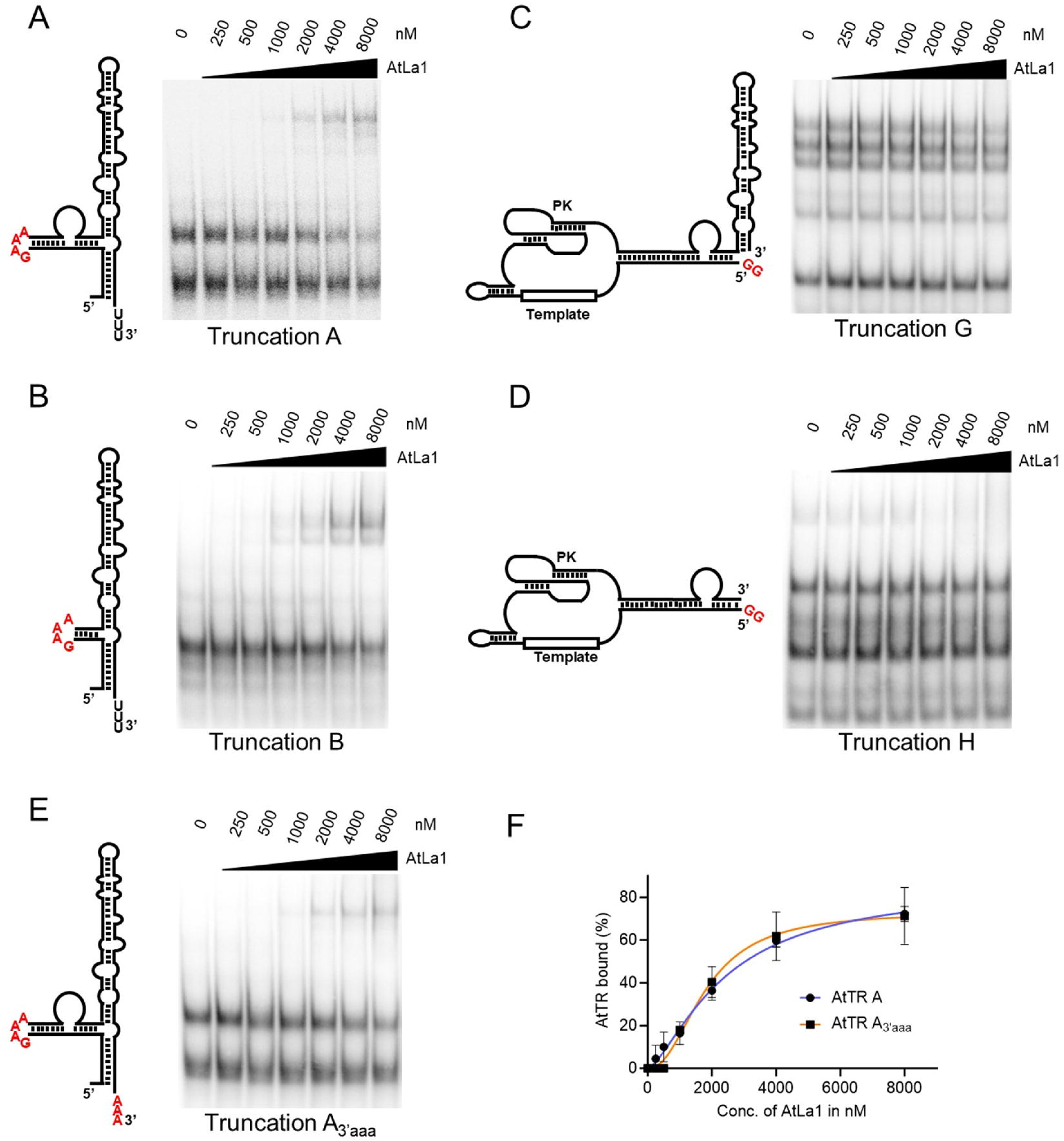
AtLa1 binds Arabidopsis telomerase RNA through the plant-specific three-way junction (TWJ). Electrophoretic mobility shift assays (EMSAs) analyzing the interaction of AtLa1_WT_ with AtTR truncation constructs. Radiolabeled RNAs were incubated with increasing concentrations of AtLa1_WT_ (0–2 μM). (A) Truncation A, lacking the pseudoknot (PK) and template region, bound AtLa1 with ∼3-fold reduced affinity (*K_d_* = 2.40 μM) compared to full-length AtTR. (B) Truncation B, which additionally lacks the bulge between P1c and P1b, bound AtLa1 with ∼2-fold reduced affinity (*K_d_*= 1.62 µM). (C, D) Truncations G and H, which retain the PK and template domains but lack the 3′ stem P1a (H additionally lacks P4–P6), failed to bind AtLa1, indicating that the TWJ is essential for productive interaction. (E) AtLa1 bound A_3’aaa_ with a dissociation constant (*K_d_*) of 1.80 μM, indicating that AtLa1 interaction is retained despite loss of the UUU-3’OH motif. (F) Quantification of EMSA data comparing truncation A and A_3’aaa_ confirms that while the UUU-3′OH residues contribute to binding, AtLa1 is not strictly dependent on this motif and likely engages additional structural elements within the TWJ.

**Fig. 6.**
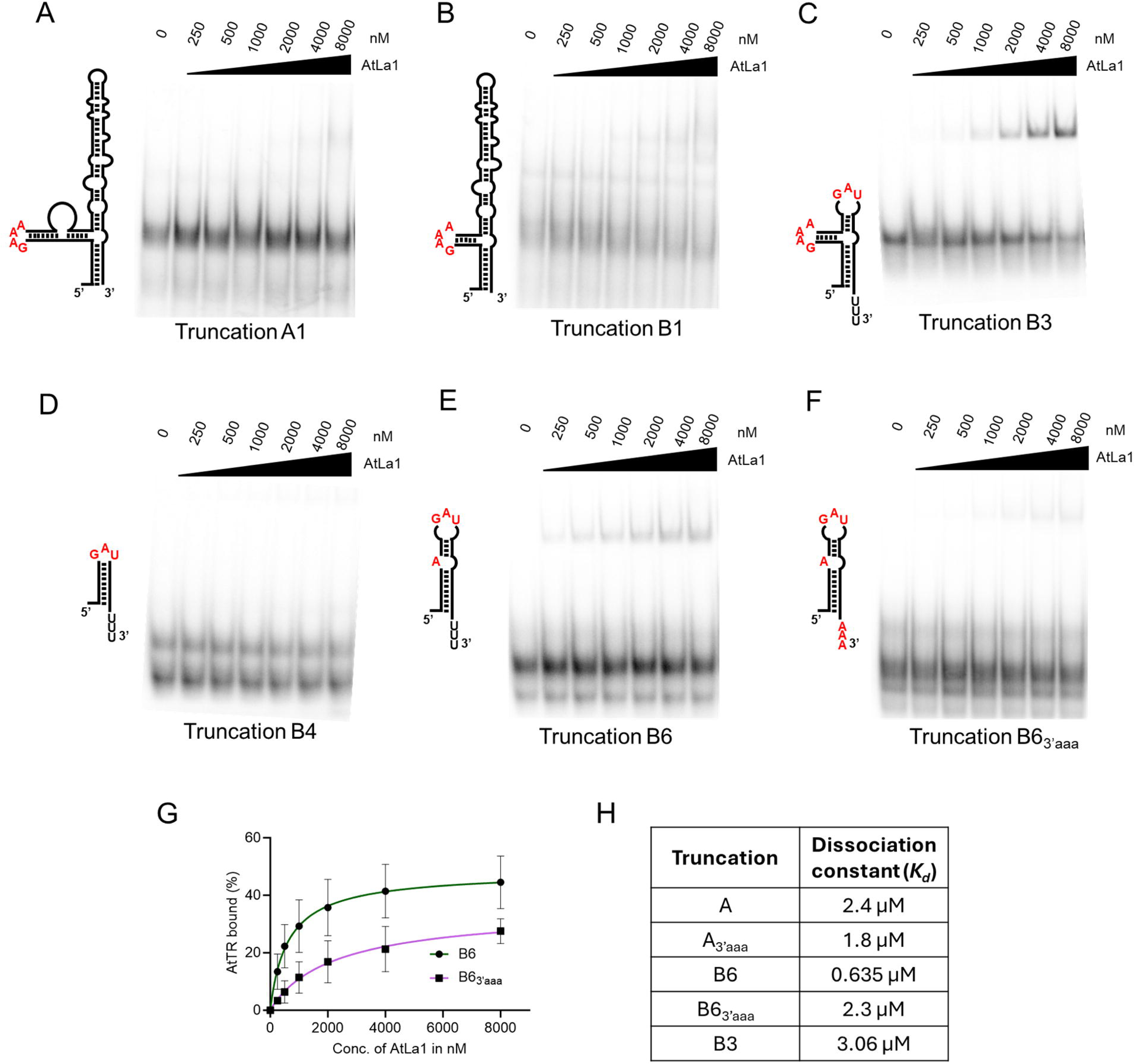
AtLa1 selectively binds to the UUU-3’OH. Electrophoretic mobility shift assays (EMSAs) analyzing binding of AtLa1 to Arabidopsis telomerase RNA (AtTR) fragments. Radiolabeled RNA constructs were incubated with increasing concentrations of recombinant AtLa1 protein (0–2 μM). (A, B) Truncations A1 (A) and B1 (B) exhibited only weak binding that was not quantifiable. (C) Truncation B3 displayed moderate binding to AtTR. (D) Truncation B4, lacking the TWJ but retaining the UUU-3′OH, showed no binding, highlighting the requirement of the TWJ for interaction. (E) Truncation B6 showed the strongest binding affinity, with a dissociation constant (*K_d_*) of 0.635 μM. (F) Replacement of the UUU-3′OH with AAA-3′OH in B6 (B6_3’aaa_) reduced the affinity by ∼four-fold, demonstrating the requirement of the U-rich tail for AtLa1 binding. (G) Binding analysis of B6 and B6_3’aaa_ reveals that the UUU-3’OH residues are essential for efficient AtLa1–AtTR interaction. (H) Summary of AtTR truncations and their binding to AtLa1.

One unique feature of AtTR is a plant-specific 3-way junction (TWJ) comprised of the P1a stem, bearing a small stretch of single-stranded RNA (SSR) consisting of a poly U stretch at RNA 3’ terminus (UUU-3’OH), the P1b stem, and the P4 stem, which appears to correspond to eCR4/5 of the human telomerase RNA (Song et al., 2019). We previously showed that the TWJ is necessary for dyskerin binding (Song et al., 2021). Notably, truncations A and B share the single-stranded part of the 3’ stem, which may serve as the site of interaction for AtLa1_WT_. Since La proteins are known to bind the UUU-3′OH of Pol III transcripts (Stefano, 1984; Gottlieb and Steitz, 1989; Wolin and Cedervall, 2002), we replaced the UUU-3’OH on truncation A with AAA-3’OH (A_3’aaa_) to determine if another domain in the RNA is contacted by AtLa1. EMSA revealed that AtLa1_WT_ interacted with A_3’aaa_ with a dissociation constant (*K_d_*) of 1.80 µM (Fig. 5E). Quantification of EMSA data comparing truncation A and A_3’aaa_ revealed that AtLa1 is not entirely dependent on the UUU-3’OH residues for interaction with AtTR, and likely engages additional structural elements within the TWJ (Fig. 5F).

We next asked if the 3’ SSR is necessary for AtLa1 binding using truncations A1 and B1, which are similar to truncations A and B, but lack the 3’ SSR (Fig. 6A and 6B). EMSA revealed very weak binding insufficient to allow quantification, and no dissociation constant could be derived using ImageJ. To further refine the AtLa1 binding site, we deleted P5 and P6 from truncation B to prepare truncation B3. Additionally, we generated truncation B4 which only has 3’ stem P1a which includes a 3’ SSR region. We found that AtLa1 retains interaction with the B3 truncation with a dissociation constant (*K_d_*) of 3.06 µM (Fig. 6C). However, the interaction with AtLa1 was completely abolished with B4 (Fig. 6D). Strikingly, removal of the P1b stem on B3 to generate truncation B6 resulted in the strongest binding affinity with a *K_d_* of 0.635 µM (Fig. 6E). When we replaced the UUU-3’OH of B6 with AAA-3’OH to create construct B6_3’aaa_, there was a four-fold decrease in the affinity (*K_d_*=2.356 µM) (Fig. 6F and 6G). Quantitative comparison of dissociation constants across truncations (Fig. 6H) highlights the enhanced affinity of B6 and the reduced binding upon disruption of either the TWJ architecture or the UUU-3′OH, underscoring the cooperative contribution of these elements to AtLa1 recognition. Taken together, these results indicate that AtLa1 engages both the TWJ and UUU-3’OH of AtTR.

### AtLa1 does not enhance the enzyme activity of telomerase RNP particles reconstituted *in vitro*

Having defined the structural determinants of AtTR binding by AtLa1, we next investigated how AtLa1 influences telomerase activity. As discussed earlier, we found decreased telomerase activity in plants depleted of AtLa1, but no evidence of a change in repeat addition processivity (Fig. 3E and 3F). Based on our data, we postulate that, AtLa1 associates with AtTR during RNP biogenesis and is replaced by dyskerin when telomerase undergoes maturation. Accordingly, we predict AtLa1 will not stimulate enzyme activity in the same way that dyskerin does. To test this hypothesis, we reconstituted telomerase *in vitro* using RRL-expressed AtTERT and AtTR (Song et al., 2019) and *E. coli* expressed and purified AtLa1. To assess telomerase activity, we employed a direct elongation assay (Song et al., 2019; Song et al., 2021) to examine enzyme activity in the presence of increasing concentrations of recombinant AtLa1. Although an increase in activity was occasionally observed, the results were inconsistent and did not correlate with AtLa1 protein concentration (Fig. 7A). In addition, when we measured the number of telomere repeats incorporated in the presence and absence of AtLa1, we found no difference (Fig. 7A), arguing that AtLa1 does not stimulate overall enzyme activity or repeat addition processivity under these conditions. Similar results were obtained when the reconstitution reactions, supplemented with either full-length AtLa1 or the truncated version, AtLa1_ΔC_, which lacks the ability to bind AtTR, were evaluated by the conventional PCR-based TRAP assay (Fig. 7B). A strikingly different result was observed when recombinant dyskerin was subjected to the same in vitro reconstitution protocol. In this case, a clear enhancement of both enzyme activity and repeat addition processivity was detected (Song et al., 2021). Taken together, these findings support our hypothesis that dyskerin and AtLa1 play distinct roles in promoting telomerase activity *in vivo*.

**Fig. 7.**
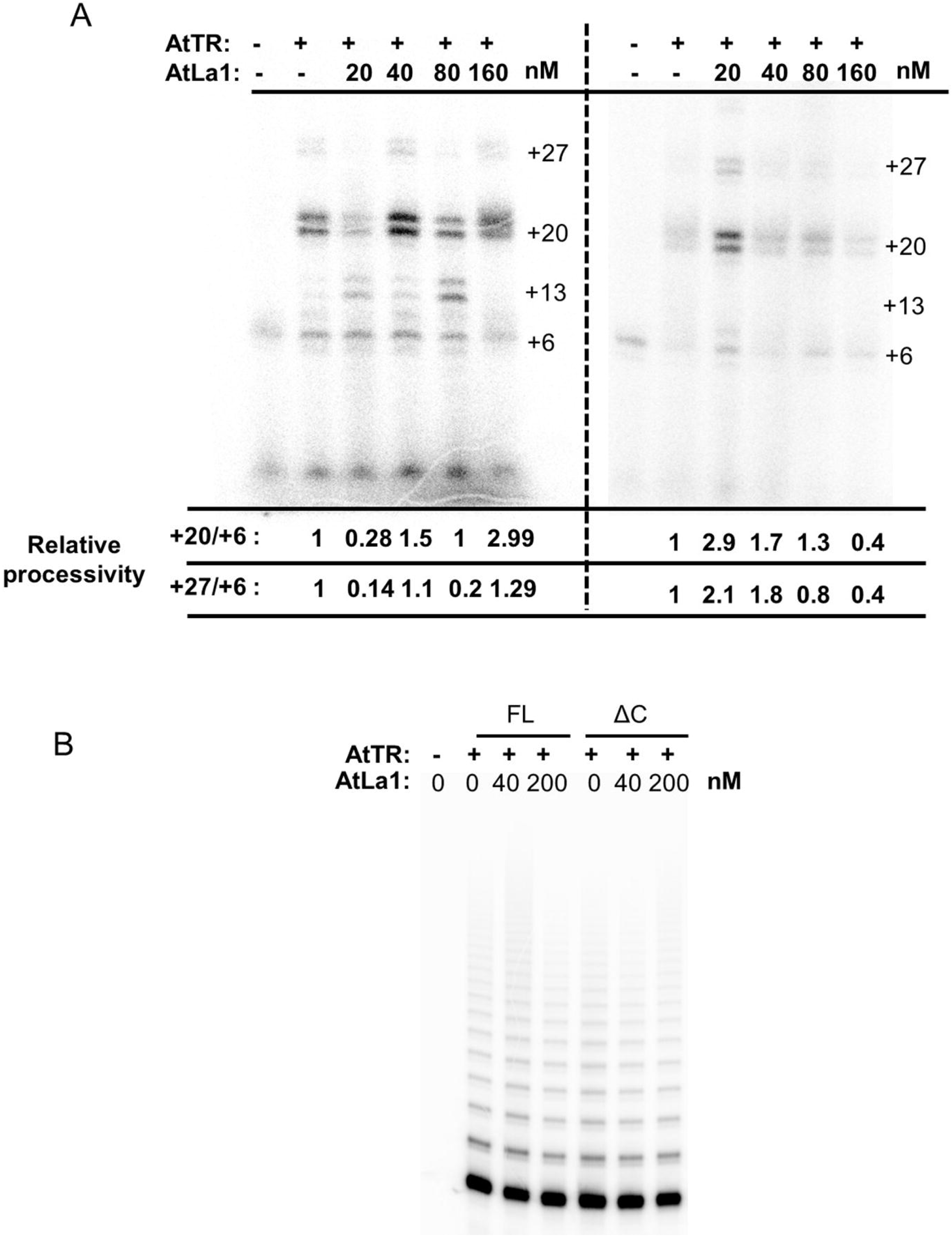
Effect of AtLa1 on Arabidopsis telomerase activity and processivity. (A) Direct telomerase primer extension assay with in vitro reconstituted Arabidopsis telomerase in the presence of increasing concentrations of recombinant AtLa1 (20–160 µM). While a slight increase in overall activity was occasionally observed, AtLa1 did not reproducibly enhance relative processivity, measured as the ratio of signal intensity between longer extension products (+27) and shorter products (+6). (B) TRAP assay. Increasing concentrations of AtLa1_WT_ and AtLa1_ΔC_ was added to the direct assay reactions. The products were used for the TRAP assay and the final products were run on the denaturing PAGE. Relative processivity was not significantly altered by either AtLa1_WT_ or AtLa1_ΔC_.

## Discussion

Telomerases display a remarkable diversity of molecular strategies to regulate biogenesis, maturation and assembly, and to ensure precise recruitment and engagement of the enzyme at chromosome termini. More recent work reveals non-canonical function in the stress response (Sharma et al., 2012; Barcenilla et al., 2023). In addition, a wide variety of telomerase accessory factors have emerged to facilitate key steps in telomerase regulation, including transcriptional control, stabilization and processing of TR, chaperoning of protein subunits, and coordination with cell cycle and DNA damage response pathways to restrict telomerase action to appropriate cellular contexts (Buchkovich and Greider, 1996; Cifuentes-Rojas and Shippen, 2012; Liu et al., 2024). Because comparatively little is known about telomerase in the plant kingdom, we employed an unbiased qMS approach to investigate whether regulatory mechanisms similar or divergent to other eukaryotic lineages exist in Arabidopsis.

The most significantly enriched protein in our MS experiment, besides tagged TERT, was AtLa1. La-related proteins (LARPs) have previously been associated with mammalian, yeast and ciliate telomerases (Aigner et al., 2000; Collopy et al., 2018; Kao et al., 2024). Tetrahymena p65 (a LARP7 family protein) induces structural changes in TR that promote its efficient binding with TERT (Aigner et al., 2000; O’Connor and Collins, 2006). Similarly, in humans and yeast LARP7 is important for telomerase enzyme activity and telomere length maintenance (Bousquet-Antonelli et al., 2009; Collopy et al., 2018; Kao et al., 2024). While no LARP-7 homologs have been identified in Arabidopsis (Xiang et al., 2024), the genuine La protein AtLa1, bearing the defining “La module,” a canonical La motif followed by an RNA Recognition Motif (RRM1) binds plant RNAs bearing a UUU-3’OH terminus, including U6 snRNA and pre-tRNA (Fleurdépine et al., 2007),

We hypothesize that AtLa1 serves as an ancestral La protein and compensates for the loss of LARP7 in the plant telomerase. In particular, we propose that AtLa1 participates in AtTR biogenesis and is retained in the initial RNP complex. Consistent with our hypothesis, AtLa1 overexpression enhances telomerase activity. Despite this rise in enzyme activity, we found no change in telomere length. However, although elevated telomerase activity levels drive telomere elongation in mammalian cells (Cristofari and Lingner, 2006), this is not the case for *A. thaliana* (Ren et al., 2004; Barcenilla et al., 2023). Telomere length is tightly regulated, and excess telomerase is apparently not utilized for telomere extension (Barcenilla et al., 2023). As predicted from our model, inducible knockdown of AtLa1 led to substantially decreased telomerase activity. Despite this decrease in activity, we saw no concomitant decrease in telomere length over the seven-day period of inhibition. Arabidopsis plants with a null mutation in *AtTERT* or *AtPOT1a* have progressive telomere shortening by 200-500 bp per generation cycle (Riha et al., 2001; Surovtseva et al., 2007). Therefore, we assume the short timeframe of AtLa1 depletion was insufficient to cause detectable telomere shortening. Overall, these findings provide strong evidence that AtLa1 is involved in telomerase biogenesis and important for maintaining telomerase function *in vivo*.

We discovered that AtLa1 interacts with AtTR *in vivo* and *in vitro*. AtLa1 binds AtTR *in vitro* with relatively low affinity (K*_d_* = 752 nM) compared to p65 and Pof8 which bind Tetrahymena TR and fission yeast TER1 with affinities of 25 nM and 30-50 nM, respectively (Akiyama et al, 2012; Singh et al., 2012). Although the biological implications of this lower affinity of AtLa1 for AtTR are unclear, it is possible that AtLa1 in its capacity as a bona-fide La protein did not co-evolve with AtTR to achieve the tighter and evolutionarily optimized RNA-protein interactions of LARP7 family members with their telomerase RNA binding partners.

We further found that AtLa1 recognizes the UUU-3’OH terminus of AtTR and the TWJ element unique to the plant telomerase RNA. Unlike canonical substrate RNAs which predominantly rely on the UUU-3’OH for AtLa1 recognition, the UUU-3’OH of AtTR seems dispensable for AtLa1 binding and can be compensated by the presence of the TWJ. How AtLa1 adapts for this plant-specific RNA element remains unclear and requires further attention. Interestingly, AtNAP57 (dyskerin) requires this same site on AtTR for binding (Song et al., 2021), prompting intriguing questions about the composition and timing of accessory factor association during telomerase biogenesis and maturation.

Unexpectedly, our purification scheme did not uncover dyskerin and AtPOT1a, the two telomerase accessory proteins previously reported to associate with TERT and TR in Arabidopsis and to promote telomerase repeat addition processivity (Kannan et al., 2008; Renfrew et al., 2014; Arora et al., 2016; Song et al., 2021). We postulate that our MS experiments with super-telomerase lines enriched for partially assembled, inactive telomerase complexes which have not yet undergone complete maturation. Because only a few particles of active telomerase are present in cells, overexpressed TERT and AtTR may have preferentially associated with factors that bind early in RNP biogenesis. We made several attempts to purify a tagged ‘native-telomerase’ so that normal stoichiometry and turnover conditions would be met by generating transgenic callus expressing TSII-tagged TERT from its native promoter. However, we were unable to successfully purify this reconstituted enzyme, likely because the abundance of native telomerase particles in cells is insufficient.

We propose that AtLa1 transiently associates with AtTR during early biogenesis to enhance RNP assembly before being displaced by dyskerin for maturation. The capacity of AtLa1 to bind AtTR, particularly to the TWJ and the 3′ U-rich single-stranded region, suggests a potential role in stabilizing immature AtTR transcripts or protecting their 3′ ends, analogous to the protective function of other La proteins on RNA Pol III products (Maraia and Intine, 2001; Naeeni et al., 2012). AtLa1 is not sufficient to promote TERT-TR assembly, enzymatic activity or processivity in telomerase *in vitro* reconstitution assays. Nevertheless, our inducible AtLa1 knockdown lines consistently showed reduced telomerase activity, but this decrease did not correlate with a decrease in the steady-state level of AtTR. Notably, the La motif and RRM1 domains of La proteins are not only implicated in RNA stabilization, but also in RNA folding and RNP assembly (Naeeni et al., 2012; Collopy et al., 2018). The Tetrahymena p65-mediated conformational remodeling of TR is reported to enhance TERT recruitment (O’Connor and Collins, 2006; Akiyama et al., 2012). Therefore, it is conceivable that AtLa1 acts during the initial steps of telomerase assembly as a chaperone for AtTR. In marked contrast to AtLa1, dyskerin enhances telomerase repeat addition processivity when added to *in vitro* reconstitution reactions containing recombinant TERT and TR (Song et al., 2021). These observations suggest that dyskerin is retained in the final RNP particle as a constitutive accessory factor in a fully active telomerase complex.

Taken together, our results provide novel insights into the biogenesis and evolution of plant telomerase. The Arabidopsis telomerase can be distinguished from other telomerase RNP complexes because it harbors a genuine La instead of a LARP7 protein. The requirement for both La1 and dyskerin highlights the cooperation of multiple RNA-binding proteins and the hybrid strategy plants employ for telomerase assembly and maturation.

## Materials and Methods

### Generation of Plant transgenic lines and growth conditions

Super-telomerase lines were created with pHSN6A01_U6AtTR_35S_gTERT construct in a *tert* mutant background (Barcenilla et al., 2023). The AtLa1 overexpression line was a generous gift (Cui et al., 2015).

### Inducible AtLa1 and AtNAp57 knockdown lines

β-estradiol inducible RNAi knockdown AtLa1 transformants were created in the wild type (Col-0) background. For plasmid construction cDNA synthesized from RNA of Col-0 with SuperScript™ III Reverse Transcriptase. 213bp of coding region of RRM1 domain of AtLa1 protein selected for construct preparation. Sense fragments were amplified with X*ho*I and *EcoR*I, and same antisense fragments were amplified with *BamH*I and *Xba*I restriction digestion sites with the help of primers, cDNA and Q5 high fidelity master mix (New England Biolabs). Both sense and antisense fragments were inserted into the hp RNAi pHANNIBAL plasmid by restriction digestion and ligation cloning. A fragment containing senseAtLa1_pdk intron_antisense AtLa1_OCS terminator was inserted into the β-estradiol inducible XVE destination plasmid pER8 with *Spe*1 and *Xho*I restriction digestion by restriction digestion and ligation cloning.

### Genetic Transformation

One-month-old wild type Col-0 plants were transformed by agrobacterium-mediated floral bud transformation with agrobacterium cells (Zhang et al., 2006). Transformed agrobacterium colonies (GV3101) were screened by plating on YEP plates supplemented with 100 µg/ml Rifampicin and 50 µg/ml Kanamycin and colony PCR. Transformed cells were suspended in a buffer (0.002 % MS, 5 % sucrose, and 0.04% silwet-77). Arabidopsis inflorescences were dipped in the solution and vacuum infiltrated for 1.5 min. Treated plants were kept horizontal and covered with plastic film and aluminum foil to retain moisture. After 24 h, plants were transferred to a standard growth chamber. T0 transformants were screened by plating on 0.5 MS supplemented with 30 μg/ml Hygromycin. T2 and T3 plants were screened for a single copy insert and homozygous lines by plating. Transformed stable lines were characterized by western blotting of seven-day-old seedlings grown 0.5 MS plate supplemented with 25 μM β-estradiol. Two stable homozygous T3 transformed inducible RNAi knockdown lines (1-7-6 and 4-3-1) were used for in vivo investigation of telomerase activity investigation. *A. thaliana* plants were transferred in 3:1 Pro-Mix Bio-fungicide and vermiculite: and grown under long-day conditions (16 h light; 8 h dark cycles).

### Quantitative Telomerase Repeat Amplification Protocol (qTRAP) and Telomerase Repeat Amplification Protocol (TRAP) assay

Total protein was extracted in buffer W (50 mM Tris-acetate, pH 7.5, 5 mM MgCl_2_, 2 mM Potassium Glutamate, 10 mM EGTA, 1.5% glycerol, v/v) with 1 mM DTT and 0.6 mM VRC. Total protein was measured by Qubit ® Protein Assay. Protein samples were diluted to a final concentration of 50 ng/µL. For qPCR, a total 525 ng protein from each sample was added to qPCR plates with 400 µM forward primer and HS dynamo SYBR green master mix for primer extension and plate was kept at 37°C for 45 min. 400µM reverse primer was added to the reaction mixture. qPCR was performed and data were analyzed. For the TRAP assay, 525 ng protein was added to reactions containing 1x primer extension buffer (50 mM Tris-OAc pH 8.0, 50mM KCl, 3mM MgCl_2_, 2mM DTT, 1mM spermidine), 0.66 µM forward primer (5’ CACTATCGACTACGCGATCAG 3’), 0.83 mM dATP, 0.83 mM dTTP and 0.83 mM dGTP. Reactions were kept at 37°C for 45 min. The reaction mixture was then subjected to Phenol/Chloroform extraction followed by overnight ethanol precipitation. After centrifugation at 15000 rpm, 4°C for 30 min, the resulting pallet was washed with 70% ethanol and resuspended in nuclease-free water. Resuspended pallet was used as the template for PCR which was also composed of GoTaq® Hot Start Colorless Master Mix (Promega), 0.4 µM forward primer (5’ CACTATCGACTACGCGATCAG 3’), 0.4 µM reverse primer (5’ CCCTAAACCCTAAACCCTAAA 3’) and 66 nM of □α-32P] dGTP – 3000Ci/mmol (PerkinElmer). After 35 cycles of PCR, the reaction was precipitated and resolved by 6% denaturing PAGE, dried, exposed and imaged on Typhoon FLA 9500 phosphorimager (GE Healthcare).

### qRT-PCR

RNA extracted from *A. thaliana* 7-day-old seedlings and one to two weeks old cell culture using the Direct-zol RNA kit (Zymo Research). cDNA synthesized from 1 µg RNA, SuperScript III reverse transcriptase (Invitrogen) and random primer mix (NEB). cDNA was five-fold diluted and added into PowerUp SyBr Green for qPCR reactions with AtTR primer provided in Supplementary and U6 snRNA is used as a reference for AtTR enrichment.

### Western blotting

Total protein was extracted from 25 mg seedling in a buffer (120 mM Tris base pH 6.8, 4% SDS,1% Bromophenol blue, 20% glycerol and 200mM DTT). Samples were lysed and denatured at 90°C for 8 min. Samples were centrifuged at 14000g for 5 min. Equal amounts of samples were analyzed by SDS PAGE with the marker run at 110V for 2 h. Protein was transferred from the gel to the activated PVDF membrane by a trans blot turbo system (Bio-Rad) and the membrane blocked in 5% milk. The membrane was incubated in 1:1000 of primary antibody (anti-AtLA1) at 4°C overnight, and after 24 h, washed thrice in the TBST buffer (TBS and 0.1 % tween-20). RuBisCO acted as a loading control. Finally, the membrane was incubated with 1:20000 IZ-rabbit HRP, visualized with an Amersham^TM^ Imager 600 and the image analyzed by Image J software.

### RIP

RNA-IP was performed with 1 g of overexpressed and wild type tissue from 7-day-old seedlings. Seedlings were crushed in liquid nitrogen and homogenized in RIP buffer (100 mM Tris-acetate, 150 mM Potassium Glutamate, 1 mM MgCl_2_, 0.5% Triton X-100, O.1% Tween-20) with 2.5 mM DTT, 20 µl/mL protein protease inhibitor (Sigma-Aldrich), and 1µl/mL RNase OUT (Thermo Fisher Scientific) and centrifuged at 5000g for 4 min at 4°C. 1/100^th^ of supernatant was kept as input, and one volume of supernatant was incubated with anti-HA tagged magnetic beads(Pierce^TM^ Anti-HA Magnetics Beads) for 2 h at 4°C and then washed with RIP buffer four times. qPCR and western blot (probed with anti-HA HRP) performed on beads and input.

### Protein expression and purification

AtLa1_WT_ (residues 1-433), AtLa1_ΔC_ (residues 1-397), AtLa1_RRM2_ (residues 298-397) and Dyskerin_ΔC_ (residues 1-439) coding DNAs were cloned into a pET28a vector with an N-terminal 6xHis-SUMO tag. The constructs were transformed into *E. coli* BL-21 DE3 Rosetta cells which were grown overnight at 30°C. Cells were resuspended in buffer (500 mM Tris-HCl, 500 mM NaCl, 5 mM imidazole, pH 7.5) and lysed by sonication. Dyskerin_ΔC_ expressing cells were lysed by a microfluidizer to obtain a higher amount of protein. The protein extract was purified on an ÄKTA go ™ protein purification system (Cytiva). HisTrap FF column (5 mL) was used to capture His-tagged proteins. The eluted fractions were assessed for size and purity by 12.5% PAGE. For AtLa1_WT_, AtLa1_ΔC,_ and AtLa1_RRM2_ fractions showing the expected protein band were subjected to SUMO protease (Sigma/Thermo) digestion, and His-tagged impurities were removed by passing the reaction through a 5 mL HisTrap FF column. The His-SUMO tag on Dyskerin_ΔC_ was not removed. Finally, AtLa1_WT_, AtLa1_ΔC_ and Dyskerin_ΔC_ were purified through Superdex® 200 Increase 10/300 and AtLa1_RRM2_ was purified through Superdex® 75 10/300 GL gel filtration columns (GE Healthcare) in buffer (50 mM Tris-Cl pH 7.3, 350 mM KCl and 2 mM DTT). Protein fractions were stored at –80°C until further use.

### Affinity purification of TERT for mass spectrometry

5g *Arabidopsis* flowers from super-telomerase plants or *tert* mutant plants (untransformed control) were ground in liquid nitrogen and homogenized in 20 ml lysis buffer (100 mM Tris-OAC pH 7.5, 150 mM KGlu, 5 mM MgCl_2_, 0.1% Triton X-100, 20 mM EGTA, 15g/L PVP, 10% glycerol, 20 μl/ml Plant protease inhibitor mixture [Sigma-Aldrich], 1 μl/ml RNaseOUT [Thermo Fisher Scientific], and 2 mM DTT). The extract was centrifuged at 4□, 13,000 rpm (>14500g) for 15 min, and the supernatant was collected. A Strep-Tactin XT 4 flow gravity column (0.2 ml) high capacity (iba: 2-5031-005) was equilibrated with 5 column volumes of equilibration buffer (100 mM Tris-OAC pH 7.5, 150 mM KGlu, 5 mM MgCl_2_, 0.1% Triton X-100, 10% glycerol, and 2 mM DTT). The supernatant was loaded onto a Strep-Tactin^®^XT 4Flow^®^ high-capacity column (IBA Lifesciences GmbH) at 4°C for 1 h. The column was washed with 2.5 ml wash buffer 1 (100 mM Tris-OAC pH 7.5, 150 mM KGlu, 5 mM MgCl_2_, 0.1% Triton X-100, 10% glycerol, 20 μL/mL plant protease inhibitor mixture [Sigma-Aldrich], 1 μl/ml RNaseOUT [Thermo Fisher Scientific], and 2 mM DTT). Samples were then washed with 2 mL wash buffer 2 (100 mM Tris-OAC pH 7.5, 150 mM KGlu, 5 mM MgCl_2_, 10% glycerol, 20 μl/ml plant protease inhibitor mixture, 1 μl/ml RNaseOUT, and 2 mM DTT). Finally, the column was washed with 0.6 ml wash buffer 3 (100 mM Tris-OAC pH 7.5, 150 mM KGlu, 5 mM MgCl_2_, 10% glycerol, and 2 mM DTT). The captured proteins were eluted with 8-column volumes of 20 mM NaOH (Strep-Tactin XT 4 Flow). 1M Bis-Tris-HCl, pH 5.8, was added to each eluted fraction to take the pH to 7.0-7.5 (final Bis-Tris concentration 30 mM). Eluted products were concentrated to 50-100 μl by a 5 kDa concentrator. (Amicon®)

### Quantitative mass spectrometry

50 µl of TFE (trifluro ethanol) was added to 50 ml affinity purified protein sample, and proteins were reduced, alkylated, and digested with trypsin as in Blank et al (PMID 32129706). Digests were filtered (10 kDa Amicon Ultra, Millipor # UFC5010BK) prior to desalting on C18 (Thermo Scientific Hypersep SpinTip # 60109-412). Peptides were dried, resuspended in 40 µl 5% acetonitrile/0.1% formic acid for LC/MS-MS using reverse phase chromatography on a Dionex Ultimate 3000 RSLC nano UHPLC system (Thermo Scientific) with a C18 trap to Acclaim C18 PepMap RSLC column (Dionex; Thermo Scientific) configuration. Peptides were eluted using a 3% to 40% gradient over 60 min with direct injection into a Thermo Orbitrap Fusion or Fusion Lumos Tribrid mass spectrometer using nano-electrospray. Data were collected using a data-dependent top-speed HCD acquisition method with full precursor ion scans (MS1) at 120,000 m/z resolution. Monoisotopic precursor selection and charge-state screening were enabled using Advanced Peak Determination (APD), with ions of charge 2-6 selected for high energy-induced dissociation (HCD) with stepped collision energy of 30% +/− 3%. Dynamic exclusion was active for ions selected once with an exclusion period of 20 seconds. All MS2 scans were centroid and collected in rapid mode. For each biological replicate data were collected for 1-3 injections. Raw MS/MS spectra were processed using Proteome Discoverer (v2.5) using the 2024 UniProt Arabidopsis thaliana reference proteome, UP000006548, a common contaminants fasta, and a PWF_Tribrid_Basic_Percolator_SequestHT workflow. Results were filtered for high confidence proteins (FDR 1%) with at least 2 peptide spectral matches (PSMs) and contaminant proteins were removed prior to statistical analysis using the edgeR algorithm implemented in Degust (https://doi.org/10.5281/zenodo.3258932). We required a minimum PSM count of 2 in at least 2 samples, and implemented the RUVr edgeR-quasi-likelihood model with TMM normalization. Mass spectrometry data are deposited in the ProteomeXchange repository (#PXD071051).

### EMSA

Telomerase RNA, its truncations, and variant RNAs were generated with an AmpliScribe^TM^ T7 High Yield Transcription Kit and purified by 6% denaturing PAGE. Purified RNAs were radiolabeled at the 5’ end with ^32^P-ATP (Revvity) using T4 Polynucleotide kinase (NEB) and PAGE purified. [^32^P] labeled RNAs were folded by heating at 98°C for 2 min and slow cooling to RT. Labeled RNAs were added to the EMSA reaction buffer (25 mM Tris-Cl pH 7.5, 50 mM KCl, 2 mM MgCl_2_, 10% glycerol, 1 mM DTT, 50 µg/ml BSA, 50 µg/ml yeast tRNA and 1µl/ 20 µl RNaseOUT [Thermo Fisher Scientific] and were incubated at 30°C for 10 min. AtLa1_WT_ was serially diluted, mixed with the RNA-containing EMSA reaction, and incubated at 30°C for 20 min. EMSA products were separated on 6% native PAGE in TBE buffer at 4°C. Gels were dried under vacuum at 80°C for 1 h before exposure to a phosphor imager screen. Data were collected on Typhoon FLA 9500 phosphorimager (GE Healthcare) and quantified using ImageJ.

### *In vitro* telomerase reconstitution

An *in vitro* telomerase reconstitution assay was performed as described previously (Song et al., 2021).

### Telomere length analysis

Telomere length measured by Primer Extension Telomere Length Analysis (PETRA) assay and Mean telomere length analysed by an online tool WALTER analysis (Lyčka et al., 2021). PETRA and WALTER analysis followed a well-established protocol from Barcenilla et al., 2023.

## Statistical Analysis

Statistical analysis is performed by GraphPad Prism 9 from MacOS and analysis is considered. We used more than three biological replicates for qTRAP analysis and results were considered statistically significant when the p-value was lower than 0.05.

## Author Contribution

C.P. and S.M. contributed equally to this work. C.P., S.M., J.S., R.H., I.K., C.C.G., E.M.M. and O.P. performed experiments. C.P., S.M., J.S., R.H., C.C.G., E.M.M., O.P. and D.E.S. analysed data. C.P., S.M., J.S., E.M.M., O.P. and D.E.S. wrote the manuscript.

## Acknowledgements

We thank members of our labs for insightful discussions. We gratefully acknowledge Dr. Margaret Glasner for providing access to her ÄKTA FPLC system, and Dr. Ping He for sharing plasmids. This work was supported in part by the National Institutes of Health (R01 GM065383 to D.E.S; R35 GM122480 to E.M.M), the National Science Foundation (MCB 2047915 to D.E.S) and the Welch Foundation (F-1515 to E.M.M).

## Data availability statement

The data that support the findings of this study are available from the corresponding author upon reasonable request.

## Short legends for supporting information

**Fig. S1.** Over-expression of AtLa1 does not alter telomere length. (A-C) Telomere length distribution in over-expression AtLa1 lines and wild type of Col-0 as measured by Primer Extension Telomere Length Analysis (PETRA) in seven-day-old seedlings. Results are shown for the telomeres on chromosome arms□1L,□2R,□and□5R.

**Fig. S2.** Transient knockdown on AtLa1 does not alter telomere length. (A) Telomerase activity (qTRAP) of wild type Col-0 is compared to β-estradiol treated Col-0. (B) Results of Telomerase activity assays (TRAP) for AtLa1 inducible RNAi knockdown lines (1-7-6 and 4-3-1) as compared to wild type Col-0. (C-E) Results of Primer Extension Telomere Length Analysis (PETRA) for telomere length distribution in seven-day-old seedling transformed an AtLa1 inducible RNAi knockdown construct (1-7-6 and 4-3-1) relative to wild type (Col-0). Plants were grown in MS plates supplemented with 25 µM β-estradiol. Results are shown for the telomeres on chromosome arms□1L,□2R,□and□5R.

**Fig. S3.** Purification of AtLa1_WT_, AtLa1_ΔC_ and AtLa1_RRM2_. SDS-PAGE analysis of collected fractions of recombinant (A) AtLa1_WT_, (B) AtLa1_ΔC_, and (C) AtLa1_RRM2_. AtLa1_WT_ and AtLa1_ΔC_ were purified through Superdex® 200 Increase 10/300 and AtLa1_RRM2_ was purified through Superdex® 75 10/300 GL gel filtration columns.

**Fig. S4.** C-terminal of AtLa1 is critical for binding with AtTR. EMSAs with AtTR truncation A reveal that AtTR cannot interact with (A) AtLa1_ΔC_ lacking c-terminal and (B) RRM2, which is conserved among LARP7 proteins.

**Table S1.** Mass spectrometry protein identifications.

