## Supplementary figures and images for "La1: an evolutionarily conserved player in the Arabidopsis telomerase complex"

### Supplementary figure 1

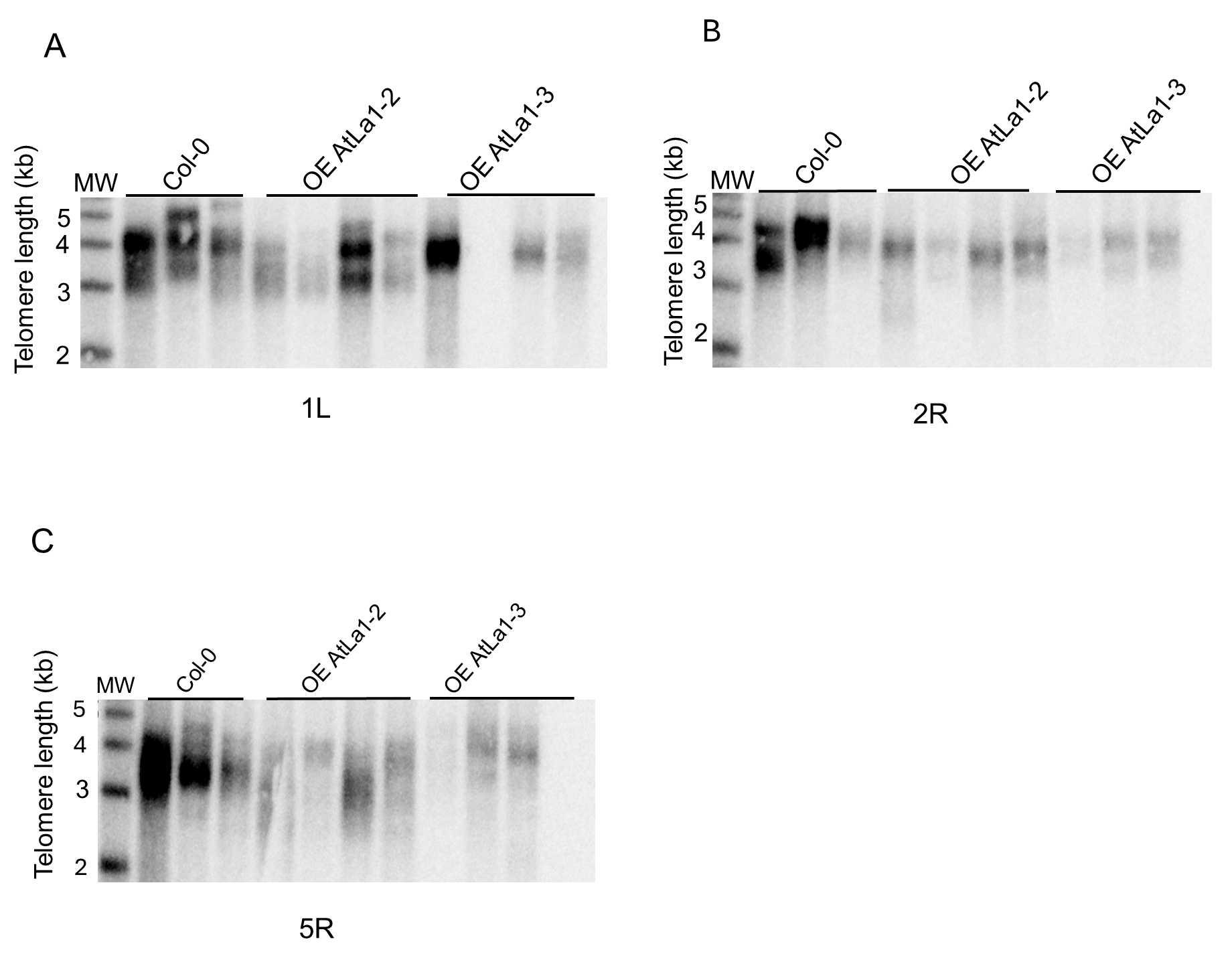

### Supplementary figure 2

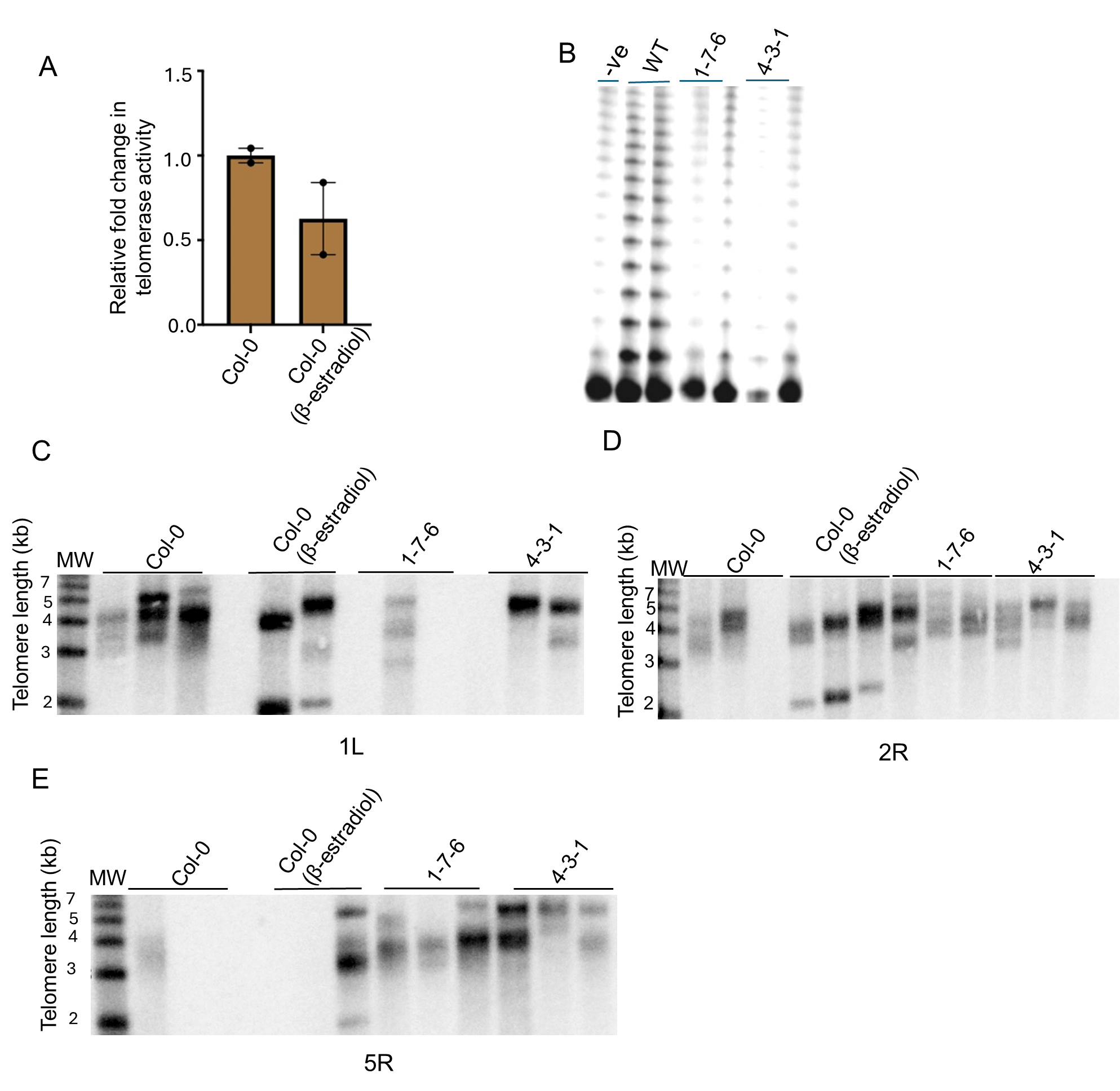

### Supplementary figure 4

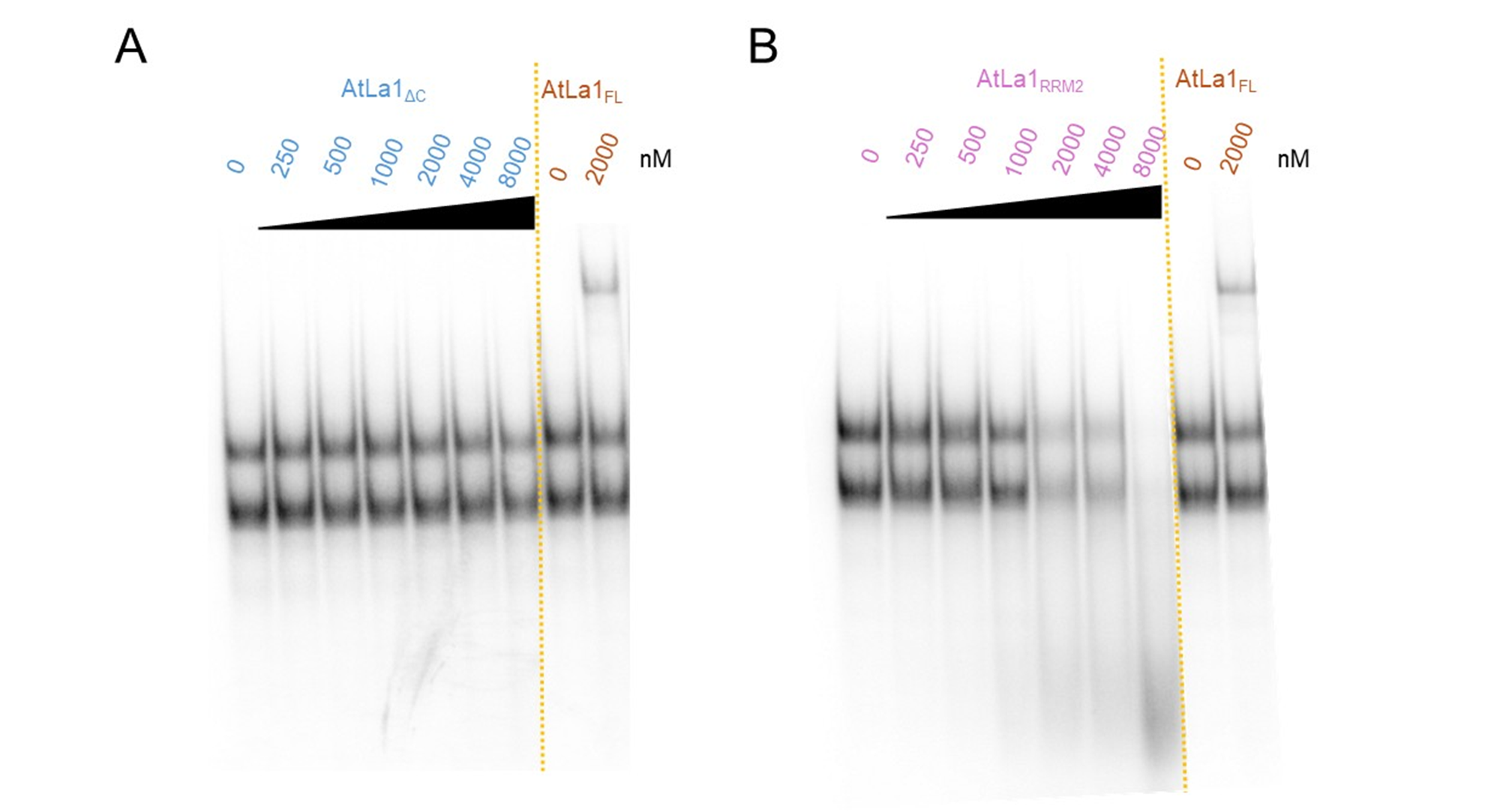
